# Müllerian mimicry in bumble bees is a transient continuum

**DOI:** 10.1101/513275

**Authors:** Briana D. Ezray, Drew C. Wham, Carrie Hill, Heather M. Hines

## Abstract

Müllerian mimicry theory states that frequency dependent selection should favour geographic convergence of harmful species onto a shared colour pattern. As such, mimetic patterns are commonly circumscribed into discrete mimicry complexes each containing a predominant phenotype. Outside a few examples in butterflies, the location of transition zones between mimicry complexes and the factors driving them has rarely been examined. To infer the patterns and processes of Müllerian mimicry, we integrate large-scale data on the geographic distribution of colour patterns of all social bumble bees across the contiguous United States and use these to quantify colour pattern mimicry using an innovative machine learning approach based on computer vision and image recognition. Our data suggests that bumble bees exhibit a manifold of similar, but imperfect colour patterns, that continuously transition across the United States, supporting the idea that mimicry is not discrete. We propose that bumble bees are mimicking a perceptual colour pattern average that is evolutionarily transient. We examine three comimicking polymorphic species, *Bombus flavifrons, B. melanopygus,* and *B. bifarius*, where active selection is driving colour pattern frequencies and determine that their colour pattern transition zones differ in location and breadth within a broad region of poor mimicry. Furthermore, we explore factors driving these differences such as mimicry selection dynamics and climate.

## Background

Mimicry has long served as an example of evolution in action, with the exceptional diversity and convergence it generates informing both microevolutionary and macroevolutionary processes [1]. Defensive mimicry occurs when there is a convergence of phenotypic qualities among organisms for the benefit of reduced predation [2]. While often defensive mimicry is considered in terms of palatable species mimicking unpalatable ones (i.e., Batesian mimicry), in Müllerian mimicry, similarly harmful, sympatric species mimic each other by converging on a shared warning signal [3]. Müllerian mimicry phenotypes, often colour patterns, are thought to be generated and sustained through frequency-dependent selection driven by predator experience: predators learn to avoid warning colours of harmful species they have previously sampled, leading to increased survival of species portraying the most abundant colour patterns [2].

It has been posited that frequency-dependent selection on a noxious lineage should promote a single, global aposematic colour pattern [4]. However, in nature, Müllerian mimicry systems follow more of a global mosaic, whereby predominant mimetic patterns differ by geographic region [5]. Such patchworks of mimetic patterns occur across Müllerian mimicry systems, with examples in the unpalatable neotropical butterflies (e.g. *Heliconius* butterflies) [6,7,8], stinging velvet ants [9,10], toxin-secreting millipedes [11], and poison frogs [12].

Several hypotheses have been proposed to explain the formation and maintenance of regional mimicry complexes. Regions may vary in the frequency of harmful phenotypes, resulting in different phenotypes being favoured in different regions [8,13]. Such variable fitness landscapes can generate alternative phenotypes even at a local scale [14,15]. The regionally favoured phenotypes will also be influenced by the ranges of predators learning the phenotypes and the extent to which the aposematic signal is memorable [16,17,18]. In addition, regional mimicry complexes can be influenced by climatic conditions, as colour patterns may vary in their thermal properties [19,20,21].

Frequency-dependent selection is argued to drive mimicry complexes to be fairly discrete, involving mimicry complexes with narrow maladaptive transition zones between them [14]. Within each of these mimicry complexes, Müllerian mimics are thought to display near perfect resemblance as predators will exert strong selection pressure against imperfect mimics [2, 22,23]. Examining features of transition zones between these complexes, such as their width [24,25,26], location, and ability to generate novel mimicry forms [27], can inform the selective processes leading to mimicry complex formation and maintenance. It has been argued, for example, that the location of these zones may lie in hotspots of species turnover, such as climatic transition zones or contact zones between historic refugia, and that partial barriers to gene flow and low population sizes in these zones may promote shifts in mimicry pattern frequencies [28]. Outside of some examples in butterflies [14,28,29,30], the properties of mimicry transition zones and the factors driving them have received little scrutiny.

Bumble bees (Apidae: genus *Bombus* Latrielle) are a particularly well-suited system to study the factors driving defensive mimicry. They display exceptionally diverse segmental colour patterning across their Holarctic range, with 427 different colour patterns across the ~250 species [21,31], a diversity explained in part by convergence and divergence onto numerous Müllerian mimicry complexes [21]. The bright aposematic colour patterns of these stinging bees have been demonstrated to be effective for predator avoidance [32,33] and geographic analysis across species revealed that their colour patterns cluster into at least 24 distinct mimicry complexes globally [21]. Mimicry also explains the exceptional polymorphism, as species that span multiple mimicry complexes tend to converge onto different mimicry patterns across their range [e.g., 15]. While examination of colour patterns at a coarse scale has been effective in defining mimicry complexes in bumble bees [21], understanding how they have evolved and are maintained requires a quantitative analysis at a fine spatial scale [5,34].

A detailed analysis of mimicry dynamics requires a reliable metric for quantification. Previous approaches analyzing colour pattern similarity by pixel [21,35] are overly sensitive to slight shifts in the location of pattern elements that may have little impact on pattern recognition by predators [21]. Coding systems that recognise relative locations of pattern elements such as colours and contrasts (e.g., two repeating stripes of yellow and black) [9,21] can be subjective, lack consideration of relative size/shape, and involve a limited number of characters. Ranks of similarity from human observers [9,36] provide metrics of mimetic fidelity, but can be sensitive to survey design.

In this study, we seek to gain a better understanding of the mimetic process by examining the extent of mimetic fidelity and the properties of mimicry transition zones in bumble bees across the United States at a fine scale. We utilise extensively georeferenced bumble bee distributional data (~ 1 in 200 Global Biodiversity Information Facility (GBIF) records is a bumble bee) and detailed documentation of their respective colour patterns to infer dynamics of mimicry. To quantify patterns of mimetic fidelity with minimal *a priori* bias, we utilised a novel machine learning-based method that calculates human perceptual distance among colour patterns. We compare transition zone dynamics among three polymorphic species, *B. flavifrons, B. melanopygus, and B. bifarius,* that shift between the Pacific and Rocky Mountains mimicry patterns by changing their abdominal segments from black to ferruginous, thus undergoing mimetic changes in parallel (Figure 2). We use these data to explore the distribution of transition zones and to determine how, apart from or in concert with frequency dependent selection, climate could potentially drive transition zone deviations.

## Materials and Methods

### Characterising general mimicry patterns

#### Mimetic fidelity of bumble bee colour patterns

To assess similarity between mimicry colour patterns, we utilised computer image recognition to determine the degree of similarity between different bumble bee colour patterns using a standardised template. A template removes effects of body size while still maintaining the “morphologically monotonous” shape of bumble bees [37], allowing us to focus only on colour pattern differences while avoiding inaccuracies from differing perspectives, light conditions, or sizes. The standardised templates used for building our colour diagrams were drawn with consideration of proportional sizes of each body segment (e.g., see Supplemental Figure 1). Colour diagrams were built for all bumble bee species occurring in the contiguous United States, excluding the rare and highly colour variable parasitic bumble bees (*Psithyrus*). In total, this included 35 bumble bee species with 63 worker patterns (queens exhibit identical or similar patterns), drawn to match colour patterns from [38]. Colours were chosen based on the colour classes used in [21] and filled using the paint bucket tool (Adobe Illustrator CC 2018) for each segmental domain. Our standardised colour palette included yellow, black, brown, white, ferruginous, or yellow-black mixed (olive) (see Supplemental Figure 1). While utilization of discrete colours will not allow for examination of colour hue or saturation variability between individuals or perceptual variability based on hue or saturation, it does allow for an improved understanding of how colour patterns as a whole, based on broad color classes, are shared among individuals and bumble bee species. Thus far, all bumble bee species have been observed to use the same pigments for these respective colours [39, 40], thus, only saturation is likely to differ substantially. Templates were exported at 256×256 pixel resolution for perceptual analyses.

**Figure 1.**
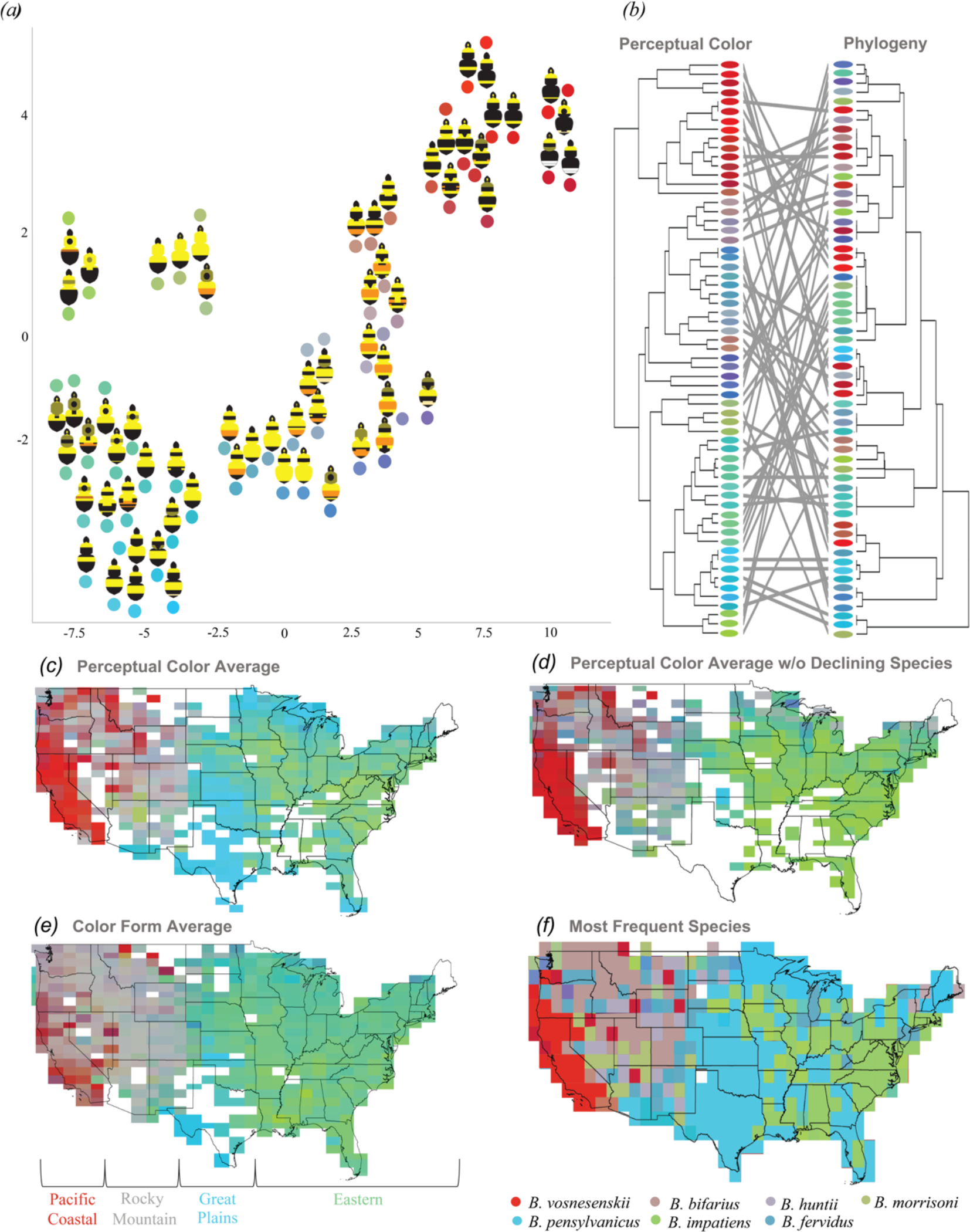
Quantification of mimetic patterns among bumble bee species of the United States. *(a)* t-SNE plot of perceptual colour pattern embedding values. *(b)* Topology of colour pattern similarity (left) compared to phylogenetic history [52] (right) with colour from the t-SNE for each taxon and tanglegram lines connecting the same taxon. *(c)* Fine-scale depiction of perceptual colour pattern frequencies calculated by averaging the t-SNE positions of all bumble bee specimens contained within each grid cell. *(d)* The same analysis, but excluding species in decline (*B. affinis*, *B. pensylvanicus/B. sonorus*, and *B. terricola/B. occidentalis). (e)* Average colour embedded score by species/colour form. *(f)* The most frequent species coloured as in the t-SNE.

**Figure 2.**
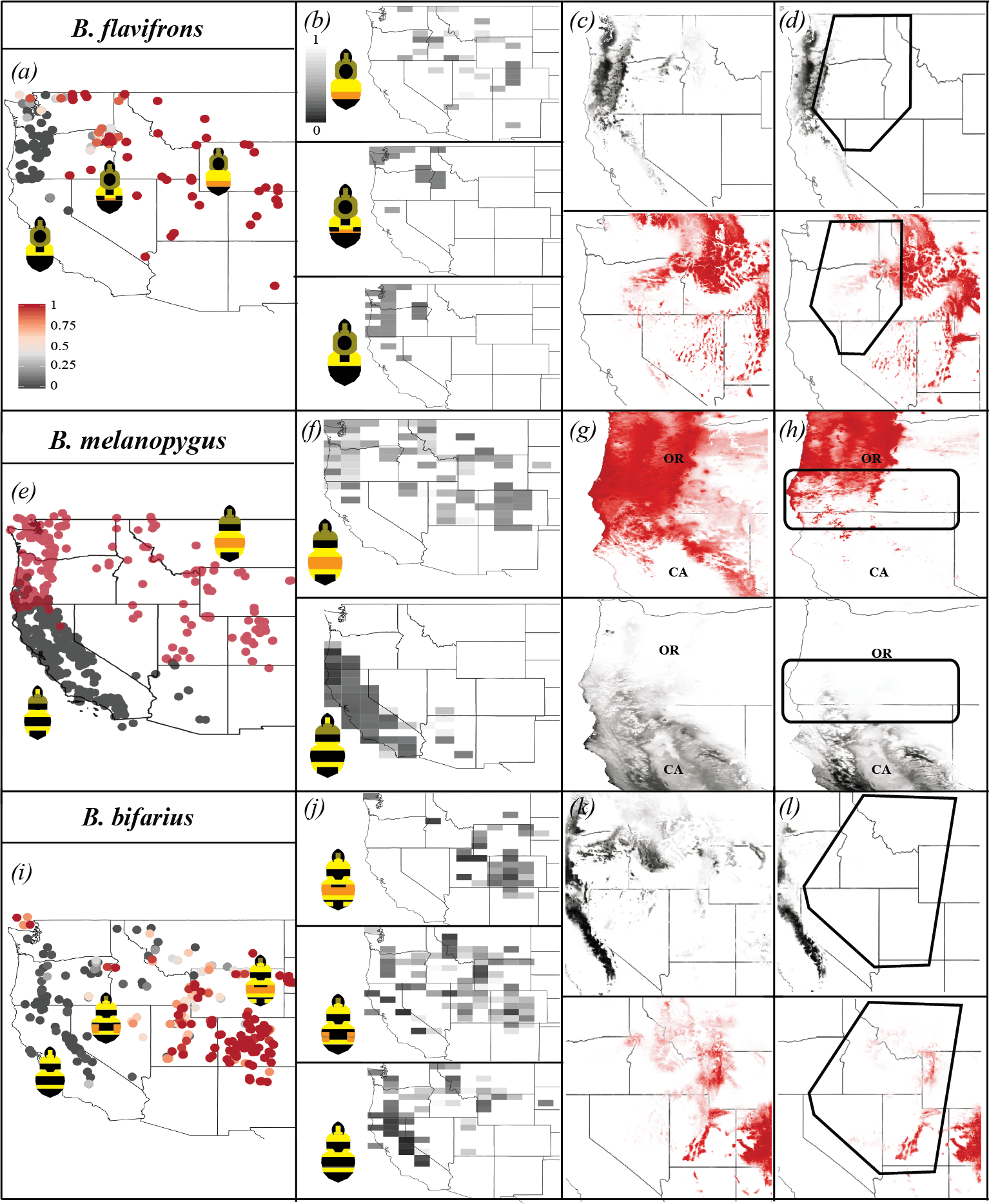
Mimetic transition zone dynamics in three polymorphic species. Left panel: point maps of colour variability and the location of colour transition zones for *B. flavifrons (a), B. melanopygus (e),* and*B. bifarius* (i), showing the extent of ferruginous, black, and intermediate variation. The scale bar represents the percent ferruginous for *B. flavifrons* and *B. bifarius,* which exhibit continuous variation. Column 2 (*b, f, j*): Regional mimetic fidelity of variable colour forms of each species, with darker colours indicating a better match to the colour pattern perceptual frequency and separate analyses run for all ferruginous, all black, and intermediate colour forms. Right columns: The impact of climatic conditions on colour form distribution with (*c,g,k*) showing niche models of full ferruginous and black colour variants modelled separately and (*d,h,l*) showing the same models when specimens from the hybrid zone (outlined regions) are excluded from the analysis.

We used an innovative machine learning approach to quantify perceptual similarity among bumble bee colour patterns. This method involves using a deep convolutional neural network (VGG-16) [41] previously trained by the Visual Geometry Group (VGG) for large-scale image recognition. The VGG-16 network uses learned visual ‘features,’ including fine scale edges, textures, and colour contrasts [41]. Distance measured in this learned feature space has been demonstrated to closely match human judgment of perceptual distance between images and significantly outperforms some of the most commonly used techniques for quantifying perceptual distance (e.g., per-pixel measures, peak signal to noise ratio) [42]. Although the predators of bumble bees remain largely unstudied, most likely they are avian [43]. While these models have not been optimized for avian perception, human visual perception has been inferred to be similar to perception of avian predators [23,35,44]. We used the feature activations of the VGG-16 network and then applied the linear rescaling of these activations from [42] to produce a distance measure between every pairwise set of bumble bee colour pattern templates. We then performed the dimensionality reduction technique, t-distributed stochastic neighbor embedding (t-SNE) [45,46], a common technique for visualizing high dimensionality distances in a 2-dimensional plot, to visualise the relative perceptual distance between all colour patterns. A colour wheel was arbitrarily applied to the t-SNE plot (Supplemental Figure 5) so that the location of each x,y coordinate and distance between x,y coordinates could be represented spatially and phylogenetically with particular colours. Distances between the colour patterns are used as a metric of mimetic fidelity with patterns closer together representing better mimics. Scripts for this analysis are available at: https://github.com/DrewWham/Perceptual_tSNE.

#### Distributional analysis of bumble bee colour patterns

To analyse geographic distributions of colour patterns, we assigned colour patterns to specimen data extracted from GBIF. We downloaded preserved specimen data for all social bumble bees found in the contiguous United States, representing a total of 160,213 records from 35 species (Fig. 1) [47]. Species pairs *B. pensylvanicus/B. sonorus* and *B. californicus/B. fervidus* were combined for this analysis given uncertain colour pattern sorting and species diagnosis between these sister species groups. Data was assessed for accuracy based on accepted distributions of species ranges [cf. 38] and localities falling well outside the documented range were removed. We confirmed this dataset to be robust for assessing relative abundance of species at a fine scale by examining raster maps of specimen abundance and species richness in 900 cells across the United States (cell size: 167 km x 88 km) constructed in R using raster [48], maps [49], and ggplot2 [50].

To determine how colour patterns cluster geographically and examine mimetic fidelity by region, we created a frequency distribution raster map depicting the perceptual average colour pattern of sympatric bumble bee species across the United States. For this we divided the United States into 900 grid cells (~167 km × 88 km) and assessed the average colour pattern among all specimens per grid cell (perceptual colour pattern frequency), a measure of overall colour pattern frequency, and among species with subspecific colour patterns considered separately (colour pattern average), a measure of colour pattern similarity among individual species. This colour pattern average removes the effect of relative frequency to assess comimicry between species.

Averages were calculated from the x,y coordinates of colour patterns from the t-SNE analysis for each grid cell. For monomorphic species, we assigned all specimens of the species the same colour pattern. For polymorphic species, we assigned colour patterns to locality data based on the literature (*B. fervidus*: [51]; *B. sylvicola*: [38,52]; *B. occidentalis*: [38,52,53]; *B. pensylvanicus*: [38]; *B. bifarius*: [54]; *B. flavifrons*: [53]) and from data obtained herein (Supplemental Figure 2). *B. rufocinctus* was excluded as it is usually rare and often misidentified due to colour pattern variability [38,52]. Grid cells containing fewer than 10 specimens (perceptual colour pattern frequency) and less than two unique colour patterns (colour pattern average) were removed from the analysis. Spatial analyses were performed in R using raster [48] and BBmisc [55].

Separate perceptual colour pattern frequencies were calculated excluding three species groups that have been extensively surveyed and studied for conservation purposes: *B. affinis*, *B. pensylvanicus*/*B. sonorus*, *B. terricola/B. occidentalis*. Abundances of these species have declined drastically in the last 10-15 years, leading to focused efforts to document historical and current records of these species. By examining patterns with and without these species, we gain understanding of how perceptual colour pattern averages change with real short-term changes in populations.

Finally, to determine if specific, abundant species influence the perceptual colour pattern frequencies, we used ArcGIS 10.3 to determine the most frequent species from GBIF data in each region (resolution: cell size: ~142 km^2^; 663 cells total).

#### Assessing convergence in warning signals

An important element of mimicry is that the shared phenotypes in a region are convergent as opposed to resulting from common ancestry [13]. Previous analyses did not consider phylogeny in assessing the degree of mimicry in species [21]. To examine the degree of convergence and how colour patterns have evolved, we created a dendrogram of perceptual colour pattern distances as calculated by the convolutional neural network using the R programs ape [56] and dendextend [57] and compared this to the distances of the bumble bee phylogenetic tree [39], including each species found in the contiguous United States, using a Mantel test [58] (R program ade4 [59], 9999 matrix permutations). To visualize the phylogenetic and perceptual colour pattern relationships, we assigned the species-specific colour pattern embedding colours to the branches of the bumble bee phylogenetic tree and the dendrogram of perceptual colour pattern distances.

### Mimetic transition zone dynamics in polymorphic species

#### Mimetic distribution and fidelity

We analyzed the fine-scale distribution of colour pattern variability within three polymorphic, co-mimetic species, *B. flavifrons, B. melanopygus*, and *B. bifarius*, using detailed phenotype data collected from approximately 300 newly collected individuals and 4,921 museum specimens (*B. flavifrons*: 268; *B. melanopygus*: 2,663; *B. bifarius*: 1,990), from seven natural history museums (Supplemental Table 1). For each individual, a microscope was used to assign colour percentages (black, ferruginous, yellow) to each section of the body, including 72 regions for females and 88 for males (Supplemental Figure 3), similar to [31]. From these data, we made distribution maps (R: ggplot2 [50], ggthemes [60]) using georeferenced locality data of the colour pattern variation most relevant to mimicry, plotting overall percentage of black vs. ferruginous colour in metasomal tergites 2 and 3 for *B. melanopygus* and *B. bifarius*, and tergite 3 in *B. flavifrons*.

To understand how mimetic fidelity varies across these species’ ranges, we assessed the degree to which the colour patterns of each polymorphic species differed from the local perceptual colour pattern frequency across their distribution. For each specimen, we estimated the distance between the t-SNE x-y coordinates for its colour pattern and the perceptual colour pattern frequency for all bumble bees located in the grid cell the specimen is contained in using the Pythagorean theorem. Distances across all specimens in a grid cell were averaged for each species and relative distances plotted.

#### Potential forces driving distributions

To examine whether mimetic transition zones in these polymorphic bumble bees tend to occur in climatic transition zones, we first performed ecological niche modeling to predict the distribution of each colour pattern based on climatic preferences. If the localities for one colour morph predict a niche distribution that encompasses the range of the other colour form that would suggest they both occupy similar climatic zones. Second, we removed localities from the region of the distribution that includes the hybrid transition zone and inferred whether climatic models alone could predict the range of each colour pattern within the hybrid zone. Niche modeling was performed in MaxEnt [61] using environmental raster layers for 19 Worldclim bioclimatic variables that account for temperature and precipitation at yearly, quarterly, and monthly time scales (1950-2000 climate, 30 seconds resolution; [62]) (Supplemental Table 2). Each environmental layer was converted to an ASCII file in R using the package raster [63] and run through 15 cross-validated replicates, with model performance assessed using the AUC statistic. The output was visualised in QGIS3 [64].

## Results

### Mimetic fidelity among bumble bees in the United States

Analysis of the similarities and differences of bumble bee colour patterns displayed in the t-SNE plot (Figure 1a) shows that bumble bee colour patterns do not form discrete clusters of similar color patterns as one might expect with mimicry, but rather bumble bee colouration falls on a continuum of perceptual colour pattern variation (Figure 1a). However, there are some distinct groupings among species that are traditionally assigned to the same mimicry zones and geographic regions. In the t-SNE plot, we see a gradual shift in colour patterns from the bottom left corner to the top right corner which reflects the transition of colour pattern morphs from east to west across the contiguous United States.

### Spatial distribution of mimicry complexes

Ample specimen data is available across grid cells throughout most of the United States (median abundance/grid cell = 74), making the GBIF data a meaningful representation of communities at this spatial scale (Supplemental Figure 4). The species richness is greatest within the Rocky Mountain Region and along the Pacific Coast, matching expectations from natural species richness, suggesting we are capturing species diversity well (Supplemental Figure 4). The lowest species richness and specimen abundance occurs in the Great Plains and southern U.S. While this represents natural abundance in the Southwest, parts of the Southeast and Great Plains are likely undersampled.

In our analysis comparing the perceptual colour pattern frequency for every grid cell within the United States, it is clear that the colour patterns of bumble bees are not similar across their range, but differ in the perceptual colour pattern frequency by geographic region. Our analysis suggests that there are four major colour pattern complexes (Figure 1c and 1d) including Pacific Coastal, Rocky Mountain, Great Plains, and Eastern mimicry regions. When we removed the species in decline – *B. affinis*, *B. pensylvanicus/B. sonorus*, and *B. occidentalis/B. terricola* – from the analysis, we found evidence for three distinct mimicry complexes in the United States, matching regions previously described as Pacific Coastal, Rocky Mountain, and Eastern [21] (Figure 1d). These declining species comprise a large portion of Great Plains faunal records, most of which are *B. pensylvanicus* (Figure 1), as indicated by the drop off in useable data when they are excluded. Given that *B. pensylvanicus* was the dominant species in historic surveys for Oklahoma [65] and Texas [66], the Great Plains mimicry complex likely represents a real historic complex rather than simply an artifact of oversampling.

Our analyses of colour pattern averages by species (Figure 1e) and the regional most frequent species (Figure 1f) allow us to assess if comimicry is occurring and if there is a single, highly abundant species driving frequency-dependent selection (Figure 1e and 1f). These analyses suggest that while the most frequent species for the Great Plains and Eastern regions are *B. pensylvanicus* and *B. impatiens* respectively, other species in this region are converging onto similar colour patterns, as the color pattern average occupies a similar colour pattern space to these species. The Rocky Mountain complex does not appear to have a single most frequent species, but rather varies between *B. bifarius* (Central Rockies), *B. huntii* (Southern Rockies), and *B. morrisoni* (S. Rockies/Desert interface) and the color pattern average occupies similar colour pattern space, supporting comimicry across species. Within the Pacific complex, while comimicry seems to occur in southern California, in northern California and Oregon there are likely many species with low abundance that are not converging onto the Pacific colour pattern and the dominant species, *B. vosnesenskii,* is likely driving the perceptual colour pattern frequency.

From this analysis, we gain improved understanding of where the predicted hybrid zones between these complexes fall. The data reveal broad, rather than narrow transition zones, between mimicry complexes with regions of high admixture and poor mimicry in between higher fidelity regions (Figure 1c). The Rocky Mountain/Pacific hybrid zone falls in eastern Oregon and Washington and is broken by the Nevada desert. The Pacific Northwest region, however, generally has a lot of mixture across the colour pattern space suggesting colour pattern diversity. The second hybrid zone lies in the transition between endemic western and eastern *Bombus* fauna as one descends the Rocky mountain foothills in the Great Plains states (Figure 1c).

### Evolution of shared warning signals

In examining bumble bee colour patterns within the United States mapped onto the phylogeny, it is apparent that similar colour patterns are scattered across distant lineages, showing little phylogenetic signal. In addition, there was no significant correlation between phylogenetic and perceptual distances (r=0.0701, p=0.1051) (Figure 1b).

### Mimetic transition zones in polymorphic species

The *B. flavifrons* transition zone aligns fairly well with that for all species (Figure 1c, 2a). *B. flavifrons* appears to exhibit relatively good fidelity outside the colour pattern transition zone, but is a poor mimic within the transition zone (Figure 2b). The primary *B. melanopygus* hybrid zone is especially narrow and shifted westward from the typical hybrid zone of all species (Figure 2e, 1c). The ferruginous and black forms are good mimics where these forms typically occur (Figure 1c), but where there are no other abundant ferruginous species, in the Pacific Northwest to northern California, this pattern is a poor mimic (Figure 2f). In *B. bifarius,* the inferred hybrid zone is much broader and more eastern shifted relative to the other polymorphic species (Figure 2i) and the standard hybrid zone (Figure 1c), with this zone transitioning from black to ferruginous with a continuum of intermediate colour patterns (Figure 2i). The fidelity of the black, ferruginous, and intermediate forms each varies with the highest fidelity observed on the far western and eastern extents of this species range. It is especially apparent that the black form is shifted further east than is optimal in the eastern portion of its range and the intermediate form sustains decent fidelity throughout its distribution suggesting that it may garner protection in both predominately black and ferruginous populations (Figure 2j).

Niche models fail to predict that colour patterns occur within each other’s range for all three polymorphic species (Figure 2c, 2g, and 2k), suggesting climatic differences between the regions of each form. When localities from each respective hybrid zone are removed, climate somewhat predicts the ferruginous colour morph distribution of each species. However, it does not predict the range of the black form, suggesting that the black forms occupy a more distinct climatic region and that each species extends beyond this climate optimum for this form.

Data files and R scripts for spatial and evolutionary analyses are available on GitHub: https://github.com/bdezray/Bumble-bee-Color-Mimicry

## Discussion

Müllerian mimicry systems exhibit exceptional diversity of colour patterns due to their colour pattern convergence within and divergence between mimicry complexes, making them model systems for understanding how phenotypes evolve through time. Several studies have examined the distributions of mimetic forms in a broad sense to reveal general patterns of mimicry [9,11,21]. Analyzing spatial patterns of mimicry at a finer scale can better reveal how mimicry complexes form and are maintained [5,34], but requires access to high resolution data. The exceptional distributional and colour pattern data of mimetic bumble bees allows for such fine-scale analyses. A major challenge, however, is to find a way to quantify colour pattern similarity with enough accuracy and precision for detailed analysis of mimetic fidelity. The use of traditional pixel-based approaches in bumble bees is error-prone [21] as colour elements often shift slightly by segment (e.g., a yellow stripe in the 4^th^ vs. 5^th^ abdominal segments would be coded completely different, but look nearly the same; Figure 1a) and binning approaches [21], where pattern contrasts are assigned to subjective colour pattern groups, forces potentially continuous patterns into discrete categories. The machine learning approach we used is modeled on human visual perceptual data obtained with the explicit purpose of extracting the most recognisable features and colour contrasts of an image to assign scores of similarity between images. These perceptual models are thus ideal for scoring perceptual colour pattern differences of mimics using image recognition criteria, enabling quantification of mimetic fidelity and distribution.

### Müllerian mimicry complexes in bumble bees are continuous and transient

Our analyses further supports mimicry in bumble bees: similar colour patterns cluster in distinct geographic zones (Figure 1c) and are convergent, as they map to very different places in the phylogeny (Figure 1b). The geographic data supports the three mimicry complexes for U.S. bumble bees (Pacific Coastal, Rocky Mountains, and Eastern) previously recognised at a coarse scale [21], as well as an additional Great Plains mimicry complex in the full analysis (Figure 1c and 1d).

Our data highlight the transience of optimal mimicry patterns. In *Heliconius* butterflies, mimicry complex boundaries were observed to shift over decades, raising the possibility that mimicry is highly evolutionarily transient in the face of frequency dependent selection [28,67,68]. In our data, the Great Plains mimicry complex is only present when species groups *B. affinis, B.pensylvanicus/B. sonorus,* and *B. terricola/B. occidentalis* are included in the analysis (Figure 1c and Figure 1d). These species have sharply declined during the last 10-15 years, thus, removing these species from the analysis demonstrates real short-term changes. The shift in inferred mimicry complexes reveals the sensitivity of mimicry optima to such changes and that ever-changing selection regimes can continuously shift mimicry patterns.

Mimicry theory states that frequency-dependent selection should pull species toward a shared more “perfect” colour pattern for each region [5], generating discrete mimicry complexes. Previous research examining the geographic clustering of similar colour patterns using a binning approach at a coarse geographic scale, ascribed bumble bees to multiple discrete mimicry groups [21]. While we recognise the same mimicry groups, we found that when examined at a fine-scale these patterns are not discrete, as bumble bees exhibit a manifold of perceptual colour patterns in colour pattern space (Figure 1a). This colour pattern continuum reflects their geographic distribution from west to east, suggesting that colour patterns tend to transition more on a gradient across geographic space as opposed to occupying discrete groups. So, how is it that bumble bees exhibit such a gradation of variable patterns with different degrees of fidelity to a regional pattern?

In nature, mimicry is often imprecise [22]. Most likely some bumble bee colour patterns are imperfect mimics of any given mimetic form. Our data shows a broad region with poor mimicry in the Pacific Northwest where mimicry zones transition. This region appears to display many different colour patterns at low frequency. In such cases, frequency-dependent selection would result in reduced directional selection favouring any given phenotype. Selection for colour pattern convergence may also be relaxed in these regions as predators may generalise colour patterns to avoid making a harmful mistake [69,70]. Some of the taxa we observed displaying patterns in between traditional mimetic patterns in our t-SNE analysis (Figure 1c) span mimicry complexes. For example, *B. mixtus* crosses the Pacific Coastal and Rocky Mountain mimicry complexes while remaining monomorphic in colour pattern. It does not directly mimic either perceptual colour pattern average, but rather, has a pattern that fits in between other colour pattern clusters (Figure 1a, Supplmentary Figure 1), thus likely acting as an imperfect mimic of both regions. Imperfect mimics partially matching both mimicry patterns in transitional zones were also observed in velvet ants [70]. Arguably, in the transition zone, an intermediate pattern between two mimicry complexes may actually be favoured, as it may best represent a perceptual average of all forms. All of these factors (reduced frequency-based selection, relaxed selection, selection favouring intermediates) could conceivably generate our observed gradient of mimicry colour patterns.

The ability of predators to recognise and remember colour patterns [71], a potential issue with fast-flying bumble bees, should also be considered. Given that some mimicry zones with nearly identical patterns exist (e.g., Southern Rockies), most likely predators can discriminate and select for more perfect mimicry. However, this could be influenced by the predator community present, flight ranges of experienced birds in the transition zone, and the ease of detecting colour patterns by habitat.

### Colour transition zones inform the evolution of mimicry

Contact, hybrid, and phylogeographic break zones tend to be shared across organisms [72,73]. Such hotspots may occur because they are midpoints between historical geographic refugia, transition zones between current habitats, and boundaries of anthropogenic change [72,73,74,75,76]. Our data support major transition zones in mimicry complexes occurring in recognised zones of faunal turnover, suggesting they are likely impacted by similar historical factors to those driving species barriers. The eastern transition zone is a common faunal transition zone [73], including for bumble bees, which exhibit nearly complete species turnover as one descends the eastern foothills of the Rocky Mountains. The transition zone between the Pacific Coastal and the Rocky Mountain mimicry complexes is sandwiched between two major mountain chains and a desert, thus supporting the hypothesis that major barriers to dispersal and gene flow could drive the distribution of contact zones [13,73,77]. There is sharing of Pacific and Rocky mountain fauna, although there are several species restricted in their distribution to the Pacific states, suggesting this transition zone naturally acts as a climatic or topographic barrier. The transition zone between the Pacific Coastal and the Rock Mountain mimicry complexes is also a very common hybrid zone in other species [73].

Polymorphic species are particularly informative for expanding our understanding of the evolutionary processes driving the diversity and distribution of colour pattern mimicry because they represent active selection on the frequency of colour pattern alleles. Our analysis examining polymorphic species revealed the locations of transitions zones to differ between *B.flavifrons*, *B. melanopygus*, and *B. bifarius*. The transition zone of *B. flavifrons* matches the general bumble bee transition zone observed between Pacific and Rocky Mountain complexes, while the transition zone in *B. melanopygus* is shifted westward and *B. bifarius* eastward. Our analysis of regional mimetic fidelity suggests that the ferruginous morph of *B. melanopygus* and black morph of *B. bifarius* extend into regions where these patterns are suboptimal (Figures 2f and 2j), thus, should be selected against over time. The transition zone of *B. melanopygus* is located in a region where many other hybrid zones occur, recognised as a contact zone generated upon glacial recession [73]. A possible historical scenario for this species is that a more widespread ancestral ferruginous form (the sister taxon *B. lapponicus/B. sylvicola* has a mostly ferruginous pattern) became isolated in southern refugia where it evolved a fixed black form to converge onto a local mimicry pattern, later to come into secondary contact in its current location. The distribution of the ferruginous form could thus be explained by the leading-edge hypothesis where species occupying northern glacial refugia were able to fill in the unoccupied space post glaciation more rapidly, therefore, preventing the northward movement of southern populations [74,75,76]. The location of intermediate patterns in *B. bifarius* falls in an area with less perfect mimicry (Figure 1c and 1d), which could enable it to be a partial mimic of both colour forms. Intermediate patterns could variably serve an advantage or at the very least be under weaker forces of selection. *B. bifarius* is also a very abundant species and, as such, whatever its pattern is in a region could become a more locally favoured pattern (it was the most abundant species in many grid cells; Figure 1f).

Colour pattern inheritance dynamics, including genetic dominance and genetic complexity and constraints, could also influence the location of mimicry transition zones. The ferruginous-black colour pattern variation observed in *B. melanopygus* is discrete (Figure 2e) and controlled by a single Mendelian gene [78], whereas *B. bifarius* exhibits a gradation of intermediate colour forms (Figure 2i) suggesting multigenic regulation. The greater complexity in gene regulation in *B. bifarius* may result in the wider hybrid zone in this species compared to the narrow hybrid zone in *B. melanopygus*, as it would need to change multiple, potentially unlinked, genes to attain the desired phenotype. Furthermore, the ferruginous morph is dominant over the black morph for *B. melanopygus*. This difference in dominance could hypothetically impact the movement of hybrid zones following the principle of dominance drive [68,79], whereby dominant alleles generate their phenotypes in both homozygous and heterozygous forms and can replace recessive alleles through frequency dependent selection. Thus, dominance drive could explain the movement and ultimately mismatch of the ferruginous form of *B. melanopygus* westward into the Pacific Coastal complex.

## Conclusions

Bumble bee taxonomy and identification has often relied on differences in their vibrant, aposematic colour patterns, resulting in the availability of exceptional documentation on colour pattern data in this system. The abundance, importance, general charisma, and ease of by-sight identification of these pollinators has also resulted in them being one of the most heavily georeferenced taxa. Utilizing this exceptional data resource and a novel application of a machine learning approach based on human visual perception, our data supports the geographic convergence of colour patterns among bumble bees as expected for Müllerian mimicry. However, counter to mimicry theory and traditional interpretation of mimicry complexes, our data suggests mimicry in bumble bees involves a continuous spectrum of colour patterns both in colour pattern space and geographically, demonstrating that mimicry is more complex and far less discrete than previously thought. This continuum may result from relaxed selection in mimetic transition zones that permits and may even favour imperfect mimicry. In addition, based on the shift in regional colour pattern optima upon removal of declining bumble bee species, it is apparent that perceptual colour pattern averages are transient over relatively short time scales. We observed the locations of mimicry transition to differ among polymorphic species, which can be explained by relaxed selection in these zones, but also historical biogeography, climate, dominance drive, and genetic complexity.

The applied machine learning approach is perhaps the most rigorous method applied to date to quantify mimetic fidelity, thus enabling fine-resolution analysis. With improved understanding of the visual properties of bumble bee pigments and the variation in these properties across species, adjustments for avian vision could be applied to digital images of bumble bees during pre-processing, prior to implementation of the convolutional neural network, to better characterize the ability of these predators to discriminate between bumble bee colour patterns.

## Acknowledgments

We acknowledge Rebecca Sommer, Luca Franzini, Sarthok Rahman, Li Tian, and Patrick Lhomme for their support in acquiring specimens. Thanks to Sydney Cameron, Jamie Strange, and Jeffrey Lozier for initial discussions. We thank Margarita Lopez-Uribe, Christina Grozinger, Erica Smithwick, Diana Cox-Foster, and Elyse McCormick for comments on the manuscript. This research was supported by NSF CAREER Grant DEB1453473 to H. Hines.

