## Supplemental Figures and Tables for "Müllerian mimicry in bumble bees is a transient continuum"

| | | | | | |
|---|---|---|---|---|---|
| *B. affinis* (af) | 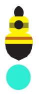 | *B. flavifrons* (fl) | 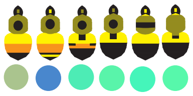 | *B. occidentalis/ terricola* (oc)/(te) | 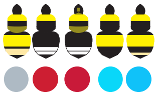 |
| *B. appositus* (ap) | 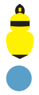 | *B. franklini* (fr) | 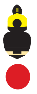 | *B. pensylvanicus/sonorus* (pe)/(so) | 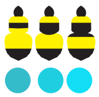 |
| *B. auricomus* (au) | 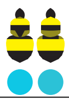 | *B. fraternus* (fa | 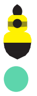 | *B. perplexus* (pr) | 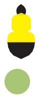 |
| *B. balteatus* (ba) | 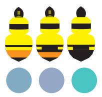 | *B. frigidus* (fi) | 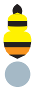 | *B. rufocinctus* (ru) | 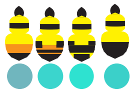 |
| *B. bifarius* (bf) | 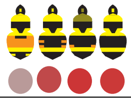 | *B. griseocollis* (gr) | 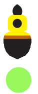 | *B. sandersoni* (sn) | 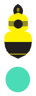 |
| *B. bimaculatus* (bi) | 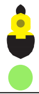 | *B. huntii* (hu) | 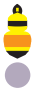 | *B. sitkensis* (si) | 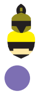 |
| *B. borealis* (bo) | 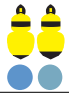 | *B. impatiens* (im) | 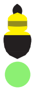 | *B. sylvicola* (sy) | 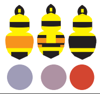 |
| *B. caliginosus* (cl) | 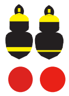 | *B. melanopygus* (me) | 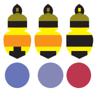 | *B. ternarius* (tr) | 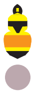 |
| *B. centralis* (ce) | 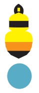 | *B. mixtus* (mi) | 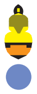 | *B. vagans* (vg) | 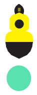 |
| *B. crotchii* (cr) | 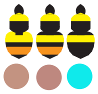 | *B. morrisoni* (mo) | 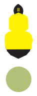 | *B. vandykei* (va) | 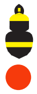 |
| *B. fervidus/ californicus* (fe)/(ca) | 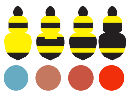 | *B. nevadensis* (ne) | 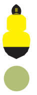 | *B. vosnesenskii* (vo) | 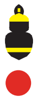 |

**Supplementary Figure 1.** Colour pattern diagrams used in perceptual and spatial analyses (Figure1) and their assigned colour embedding from the t-SNE analysis by species.

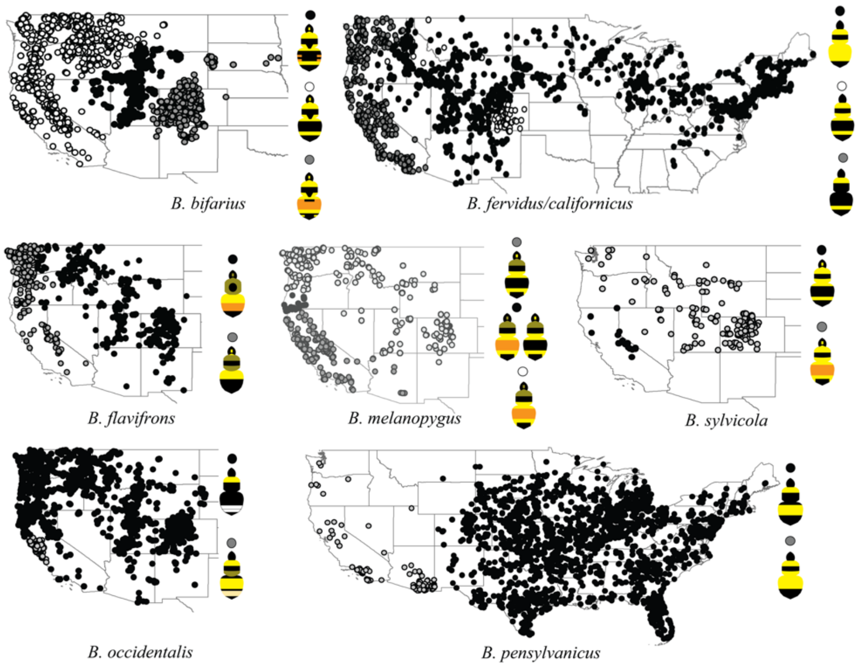


**Supplemental Figure 2.** Assignment of colour forms of polymorphic species for analysis of perceptual colour averages, assigned based on literature references of colour distribution and the data obtained for this study.

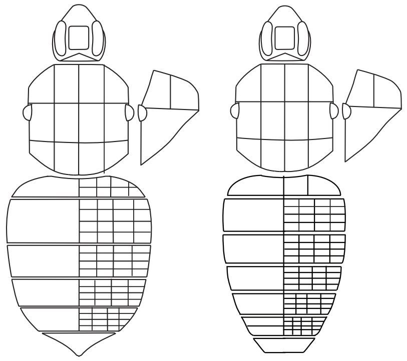

**Supplemental Figure 3.** Bumble bee template used to phenotype and quantify colour patterns of *B. flavifrons*, *B. melanopygus*, and *B. bifarius*. Colour data from the second and third metasomal segments were used for this study.

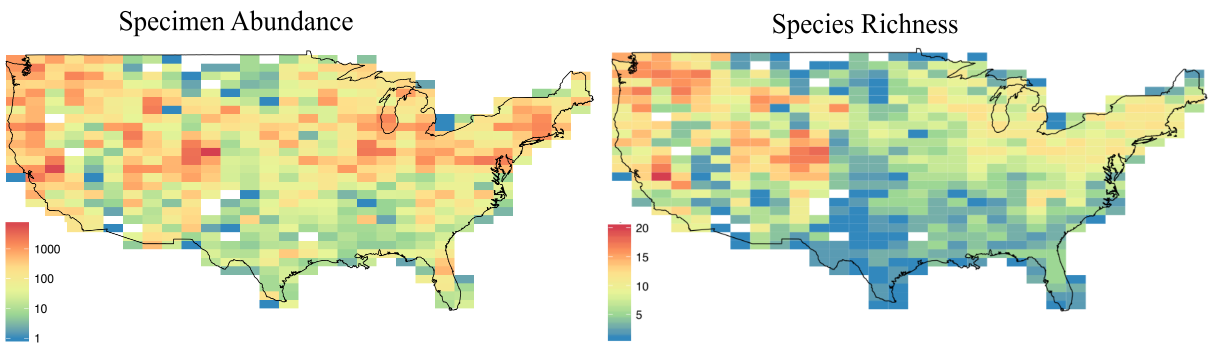


**Supplemental Figure 4.** Species abundance and species richness of bumble bee GBIF data utilized in our analyses.

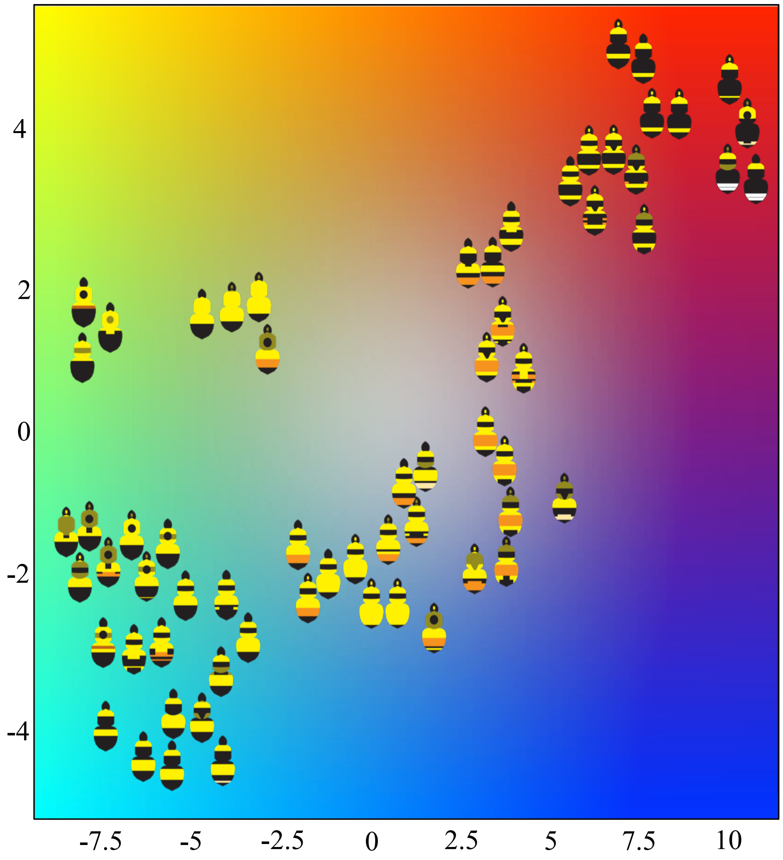


**Supplemental Figure 5.** The colour wheel utilized to extract colour coding for each x,y t-SNE embedding value.

| |
|---|
| American Museum of Natural History |
| Smithsonian Institution National Museum of Natural History |
| Peabody Museum of Natural History at Yale University |
| Essig Museum at University of California, Berkeley |
| Bohart Museum at University of California, Davis |
| The Oregon State University Arthropod Collection |
| The USDA Bee Systematics and Biology Laboratory at Utah State University |

**Supplemental Table 1.** Natural history museums visited to document colour phenotypes of polymorphic species: *B. flavifrons*, *B. melanopygus*, *B. bifarius*.

| | |
|---|---|
| 1 | annual mean temperature |
| 2 | mean diurnal range |
| 3 | isothermality |
| 4 | temperature seasonality |
| 5 | maximum temperature of the warmest month |
| 6 | minimum temperature of the coldest month |
| 7 | temperature annual range |
| 8 | mean temperature of the wettest quarter |
| 9 | mean temperature of the driest quarter |
| 10 | mean temperature of warmest quarter |
| 11 | mean temperature of the coldest quarter |
| 12 | annual precipitation |
| 13 | precipitation of the wettest month |
| 14 | precipitation of the driest month |
| 15 | precipitation seasonality |
| 16 | precipitation of the wettest quarter |
| 17 | precipitation of the driest quarter |
| 18 | precipitation of the warmest quarter |
| 19 | precipitation of the coldest quarter |

**Supplemental Table 2.** Bioclimatic variables utilised in all MaxEnt climatic niche analyses.
